# Endoplasmic-Reticulum stress controls PIN-LIKES abundance and thereby growth adaptation

**DOI:** 10.1101/2023.05.12.540474

**Authors:** Sascha Waidmann, Chloe Beziat, Jonathan Ferreira Da Silva Santos, Elena Feraru, Mugurel I. Feraru, Lin Sun, Seinab Noura, Yohann Boutté, Jürgen Kleine-Vehn

## Abstract

Extreme environmental conditions eventually limit plant growth (1, 2). Here we reveal an unprecedented mechanism that enables multiple external cues to get integrated into auxin-dependent growth programs in *Arabidopsis thaliana*. Our forward genetics approach on dark grown hypocotyls uncovered that an imbalance in membrane lipids enhances the protein abundance of PIN-LIKES (PILS) (3–5) auxin transport facilitators at the endoplasmic reticulum (ER), which thereby limits nuclear auxin signaling and growth rates. We show that this subcellular response relates to ER stress signaling, which directly impacts on PILS protein turnover in a tissue-dependent manner. This mechanism allows PILS proteins to integrate environmental input with phytohormone auxin signaling, contributing to stress-induced growth adaptation in plants.

---

Plants shape their architecture by constantly integrating environmental information into their developmental program. The phytohormone auxin is a coordinative factor between internal and external signals and provides flexibility to plant growth. Auxin is perceived in the nucleus via the TIR1/AFB family of F-box proteins, which contributes to genomic as well as non-genomic responses (6). The tissue distribution of auxin depends on a complex interplay of auxin metabolism and transport (7). The canonical PIN-FORMED (PIN) auxin efflux carriers are active at the plasma membrane and of particular developmental importance because they determine the direction of intercellular auxin transport and thereby the differential tissue distribution of auxin (8). In contrast, non-canonical PINs partially remain at the ER membrane (8). Compared to intercellular transport, the intracellular compartmentalization of auxin and its physiological roles are less well understood.

The PIN-LIKES (PILS) are predicted to be structurally similar to PINs, but are evolutionary distinct intracellular auxin transport facilitators that are fully retained at the ER (3–5). PILS proteins control the nuclear abundance and signaling of auxin (3–5), presumably by a compartmentalization-based reduction of auxin diffusion into the nucleus. External cues, such as light and temperature, define the protein abundance of PILS proteins and thereby tailor auxin-dependent organ growth rates to the underlying environmental conditions (4, 5, 9). The posttranslational control of PILS proteins can overturn the transcriptional control of *PILS* genes (5). This proposes a particular developmental importance for the control of PILS turnover, but very little is known about cellular mechanisms that integrate external cues by defining PILS protein abundance (10).

## Results and Discussion

To shed light on the control of PILS5 protein levels, we have used constitutive *p35S::PILS5-GFP* (*PILS5-GFP*^*OX*^) expression and its growth repressive effects in *Arabidopsis thaliana* (3). Here we have used the dark-induced elongation of hypocotyls (3), which is also a physiologically important response because it lifts photosynthetic organs through the soil. Using this growth model, we have conducted a non-saturated, PILS5 enhancer screen, isolating mutants that presumably impact on PILS protein abundance in a posttranslational manner (Figure 1A, B (9)). The here identified *imperial pils2* (*imp2*) mutant in the *p35S::PILS5-GFP* background (*imp2*; *PILS5-GFP*^*OX*^) displayed an increase in PILS5-GFP abundance, correlating with reduced dark-grown hypocotyl elongation (Figure 1A-C). Rough mapping and next-generation sequencing of *imp2* identified a mutation in the gene coding for the *CHOLINE TRANSPORTER-LIKE1* (*CTL1/CHER1*) (Figure 1D, E; SFigure 1A). Outcrossed *imp2* mutants in *Col-0* wild-type background were not distinguishable from the *cher1-4* loss-of-function mutants (SFigure 1B). Moreover, the overexpression of PILS5-GFP in the *cher1-4* allele induced growth retardation in dark-grown hypocotyls, being reminiscent of *imp2; PILS5-GFP*^*OX*^ mutants (Figure 1D). In addition, the expression of *pCHER1::CHER1-YFP* in *imp2; PILS5-GFP*^*OX*^ rescued the growth retardation phenotype back to the level of *PILS5* overexpressors (Figure 1E). This set of data suggests that the mutation in *CHER1* causes the defects observed in *imp2*. Accordingly, hereafter *imp2* refers to the here identified point mutation in *CHER1*, which at least partially disrupts the function of *CHER1*. Notably, *cher1-4* and *imp2* mutants as well as PILS5 overexpressors show also shorter main root growth (SFigure 1C). In contrast to hypocotyls, the repression of growth was not additive in *imp2*; *PILS5-GFP*^*OX*^ mutant roots (SFigure 1C), suggesting distinct mode of action in roots and dark grown hypocotyls.

**Figure 1:**
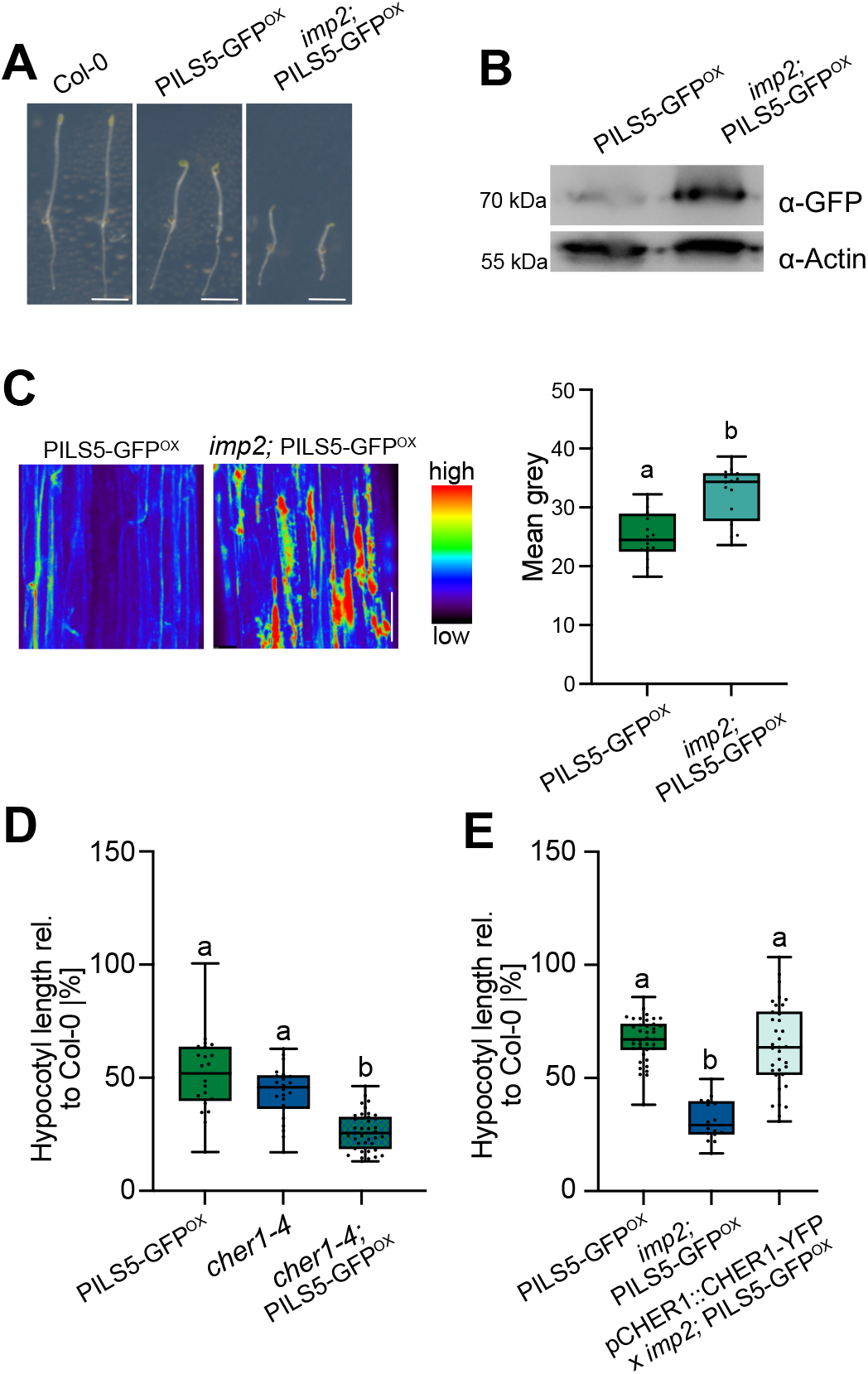
*imp2* is defective in *CHER1*. **A**, Representative images of 4-days-old dark-grown seedlings of *Col-0* wild-type, PILS5-GFP^OX^ (p35S::PILS5-GFP) and *imp2* (in the PILS5-GFP^OX^ background) grown on ½ MS. Scale bars, 5 mm. **B**, Immunoblot of PILS5-GFP in 3-days-old dark-grown PILS5-GFP^OX^ and *imp2;*PILS5-GFP^OX^ seedlings. α-Actin antibody was used for normalization. **C**, Representative images and quantifications of PILS5-GFP signal in 3-day old dark-grown PILS5-GFP^OX^ and *imp2;* PILS5-GFP^OX^ seedlings. Scale bars, 50 μm. n = 16, Student’s t-test (b: P < 0.0001). **D**, Relative hypocotyl length of 4-days-old dark-grown PILS5-GFP^OX^, cher1-4, and cher1-4; PILS5-GFP^OX^ seedlings compared to *Col-0* wild-type. n = 22-38, one-way ANOVA followed by Tukey’s multiple comparison test (b: P < 0.0001). **E**, Relative hypocotyl length of 4-days-old dark-grown PILS5-GFP^OX^, imp2; PILS5-GFP^OX^, and imp2; PILS5-GFP^OX^ complemented with pCHER1::CHER1-YFP seedlings compared to *Col-0* wild-type. n = 22-38, one-way ANOVA followed by Tukey’s multiple comparison test (b: P < 0.0001). In all panels boxplots: Box limits represent 25th percentile and 75th percentile; horizontal line represents median. Whiskers display min. to max. values. Representative experiments are shown and all experiments were repeated at least three times.

*CHER1* contributes to multiple aspects, including vascular patterning (11), ion homeostasis (12), as well as differential growth control during apical hook development (13). Most of the pleiotropic phenotypes of *cher1* mutants relate to altered levels of phosphatidylcholines in cellular membranes (reviewed in (14)). In agreement, *imp2* and *cher1-4* mutants displayed the expected alterations in phospholipid content and these defects were not modified by ectopic *PILS5* expression (SFigure 1D).

Accordingly, the imbalance in membrane lipids may impact PILS protein abundance. This assumption was further supported by the usage of the ceramide inhibitor fumonisin B1 (FB1), which disrupts sphingolipid biosynthesis at the ER (15–18). FB1 applications similarly increased PILS5-GFP abundance (Figure 2A) and also caused the enhancement of PILS5-induced growth repression in dark-grown hypocotyls (Figure 2B). Notably, FB1 treatment of PILS5-GFP^OX^ seedlings phenocopied the *imp2;* PILS5-GFP^OX^ mutant (Figure 2A, B). On the other hand, FB1 application did neither enhance the PILS5-GFP abundance nor the hypocotyl growth phenotype in *imp2;* PILS5-GFP^OX^ mutants (Figure 2A, B). We accordingly conclude that an imbalance in membrane lipids defines PILS protein abundance and growth.

**Figure 2:**
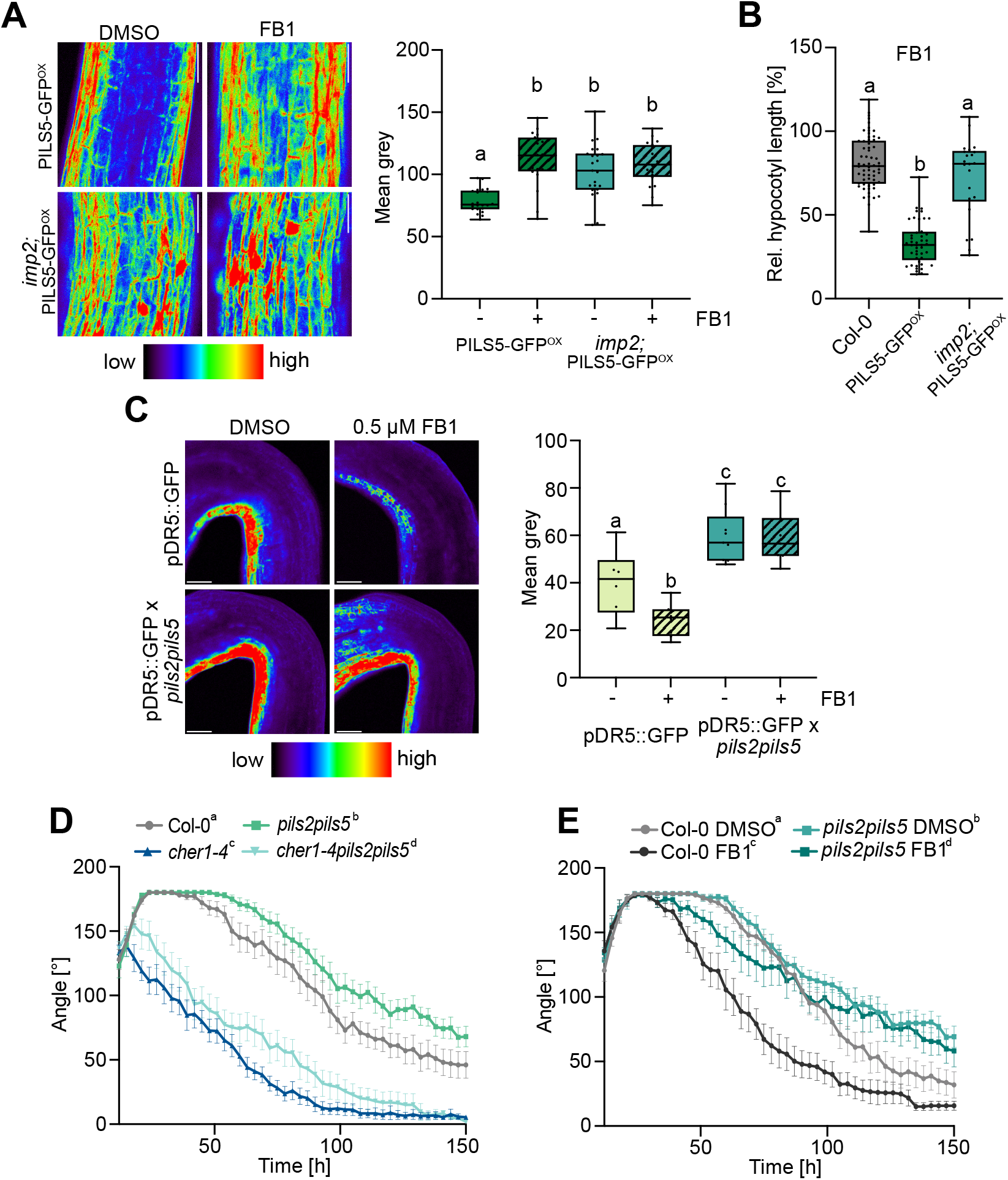
Imbalance in membrane lipids affect PILS abundance. **A**, Representative images and quantifications of PILS5-GFP signal in 3-day old dark-grown PILS5-GFP^OX^ and *imp2;* PILS5-GFP^OX^ seedlings. Seedlings were grown on DMSO (solvent control) or 0.5 μM FB1 (fumonisin B1) containing ½ MS medium. Scale bars, 50 μm. n = 20, two-way ANOVA followed by Tukey’s multiple comparison test (PILS5-GFP^OX^ DMSO vs. FB1 and PILS5-GFP^OX^ DMSO vs. *imp2*; PILS5-GFP^OX^ FB1, b: P < 0.0001; PILS5-GFP^OX^ DMSO vs. *imp2*; PILS5-GFP^OX^ FB1, b: P < 0.001). **B**, Relative hypocotyl length of 4-days-old dark-grown *Col-0* wild-type, PILS5-GFP^OX^ and *imp2;* PILS5-GFP^OX^ seedlings. Seedlings were grown on DMSO or 0.5 μM FB1 and relative hypocotyl length was calculated. n = 22-60, one-way ANOVA followed by Tukey’s multiple comparison test (b: P < 0.0001). **C**, Representative images and quantifications of pDR5::GFP signal in 4-day old dark-grown seedlings. Seedlings were grown on DMSO or 0.5 μM FB1 containing solid medium. Scale bars, 50 μm. n = 6-8, two-way ANOVA followed by Tukey’s multiple comparison test (b, c: P < 0.0001). **D** and **E**, Kinetics of apical hook opening (D) of *Col-0* wild-type, *pils2 pils5, cher1-4, pils2 pils5 cher1-4* or *Col-0* wild-type and (E) *pils2 pils5* germinated on ½ MS media supplemented with solvent control DMSO or 0.5 μM FB1. n ≥ 12, statistical significance was evaluated by non-linear regression and a subsequent extra sum of squares F test. End of maintenance phase (*X*_*0*_) and speed of opening (*K*) were compared to *Col-0* wild-type (D) or DMSO control € (b, c, d: P < 0.0001). In all panels boxplots: Box limits represent 25th percentile and 75th percentile; horizontal line represents median. Whiskers display min. to max. values. Representative experiments are shown and all experiments were repeated at least three times.

*cher1* mutants show severe growth repression in roots and shoots, but in contrast accelerated growth during apical hook opening, correlating with reduced auxin signaling rates at the concave (inner) side of the apical hook (13). In agreement, FB1 application reduced auxin signaling at the inner side of apical hooks (Figure 2C), largely phenocopying *cher1* mutants (13). This finding suggests that an alteration in membrane lipid composition affects auxin signaling in apical hooks. *PILS2 and PILS5* redundantly contribute to apical hook opening kinetics, by reducing auxin signaling at the inner side of apical hooks (4). Correlating with its effect on PILS5 protein abundance, FB1-induced repression of nuclear auxin signaling was reduced in *pils2 pils5* double mutants (Figure 2C). This finding suggests that an imbalance in membrane lipids affects auxin signaling in a PILS-dependent manner.

The PILS-induced repression of auxin signaling initiates apical hook opening (4). In agreement with an increase in PILS levels and reduced auxin output signaling, *cher1* mutants (Figure 2D; SFigure 2A (13)), as well as FB1 application showed strongly accelerated apical hook opening in the dark (Figure 2E; SFigure 2B), which is reminiscent to *PILS5* overexpression ((4, 13) SFigure 2E). Notably, FB1 treatments as well as the *cher1*-induced defects in apical hook opening were partly alleviated in *pils2 pils5* mutants (Figure 2D, E).

We thus conclude that the interference with lipid homeostasis affects PILS protein abundance at the ER, thereby contributing to auxin-dependent growth regulation.

The imbalance in membrane lipid composition did not only affect PILS protein abundance but caused the ectopic accumulation of PILS protein-containing ER structures, which likely signifies a cellular stress response at the ER (Figure 1C, SFigure 2C). In accordance, defects in lipid metabolism, including fatty acid desaturation and phosphatidylcholine metabolism, is a cellular disturbance that causes ER stress in fungal, animal and plant cells (19–25). In line with the published findings, *imp2* mutants showed transcriptional activation of ER-stress reporters (SFigure 2D), also designated as Unfolded Protein Response (UPR) genes. We, hence, tested if in fact ER stress affects the PILS5 protein abundance, using commonly used elicitors of ER stress, such as salt and tunicamycin (TM) treatments. Salt stress eventually limits biochemical processes, which lead among others to broad stress responses at the ER (26). TM is a specific inhibitor of N-linked glycosylation, thereby interfering more specifically with protein folding and consequently inducing ER stress (27). Salt as well as TM applications strongly upregulated PILS5 proteins (Figure 3A, B; SFigure 3A, B). This set of data indicates that ER-stress-inducing conditions, including imbalance in membrane lipids, salt stress, and unfolded proteins, lead to the upregulation of PILS5 proteins.

**Figure 3:**
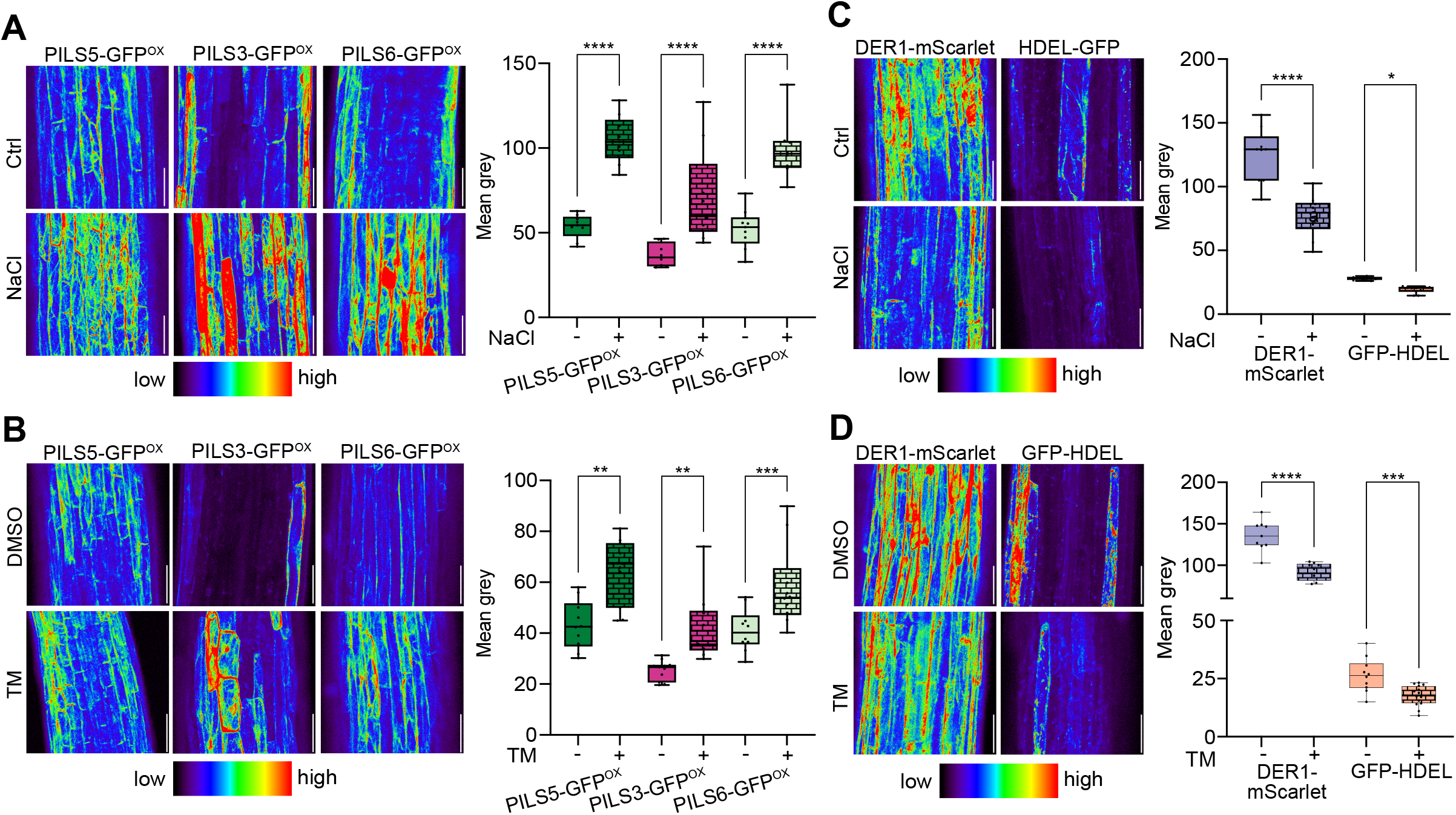
ER-stress-inducing conditions stabilize PILS protein levels. **A** and **B**, Representative images and quantifications of PILS5-GFP^OX^, PILS3-GFP^OX^ and PILS6-GFP^OX^ signal in 3-days-old dark-grown seedlings. Seedlings were grown on ½ MS and treated with or without 75 mM NaCl (A), DMSO (solvent control) or 0.5 μg/ml TM (tunicamycin) (B) in liquid ½ MS for 4h. Scale bars, 50 μm. n = 8-14, Student’s t-test between ctrl. and treatment (**P < 0.01, ***P < 0.001, **** P < 0.0001). **C** and **D**, Representative images and quantifications of p35S::DER1-mScarlet and p35S::GFP-HDEL signal in 3-days-old dark-grown seedlings. Seedlings were grown on ½ MS and treated with or without 75 mM NaCl (C) or DMSO or 0.5 μg/ml TM (D) in liquid ½ MS for 4h. Scale bars, 50 μm. n = 8-14, Student’s t-test between ctrl. and treatment (**P < 0.01, ***P < 0.001, **** P < 0.0001). Representative experiments are shown and all experiments were repeated at least three times.

Subsequently, we tested if this posttranslational effect is specific to PILS5. We observed that seedlings constitutively expressing GFP-PILS3 or PILS6-GFP showed a similar ER-stress-induced upregulation (Figure 3A, B; SFigure 3A, B). Next, we addressed whether ER stress has a general impact on ER-localized proteins. ER-stress did not increase, but contrary reduced the abundance of the ER luminal GFP-HDEL and transmembrane ER-marker DERLIN1 (DER1)-mScarlet (Figure 3C, D; SFigure 3A, B).

This set of data suggests that ER-stress-inducing conditions exert a specific effect on PILS proteins in dark-grown hypocotyls.

To assess if this response is possibly indirect, we addressed the response kinetics of ER stress-induced PILS protein abundance in dark-grown hypocotyls. Salt, as well as TM, increased p35S::GFP-PILS3 abundance within 1 hour (SFigure 4A, B), suggesting a rather direct effect of ER-stress on the posttranslational control of PILS protein abundance. Notably, we also observed a similar response for functional *pPILS3::PILS3-GFP* (4) in the *pils3-1* mutant background (Figure 4A,B; SFigure 4C, D), suggesting that ER stress also affects physiologically relevant protein levels of PILS3.

**Figure 4:**
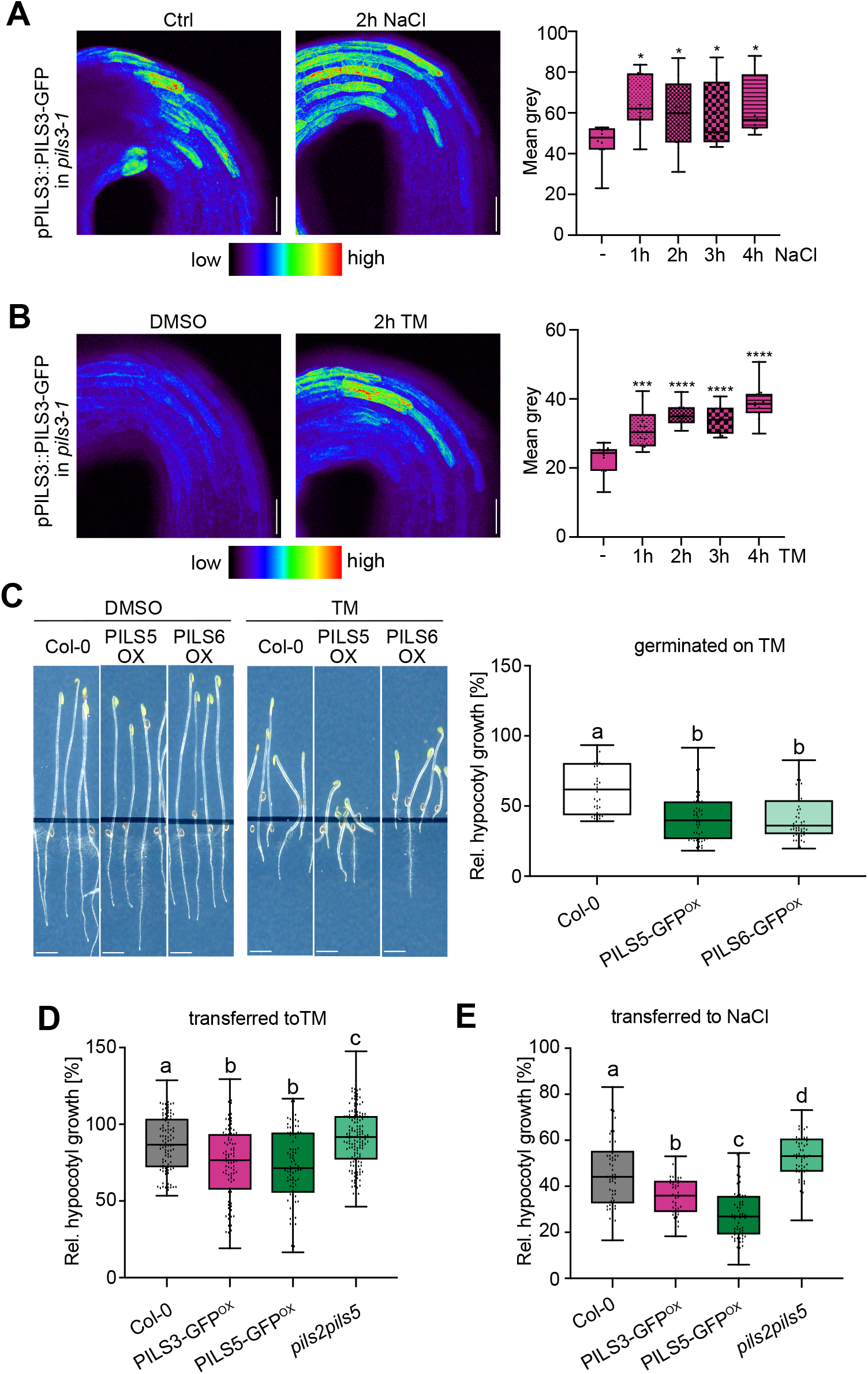
ER-stress defines PILS-dependent growth **A-B**, Representative images and quantifications of pPILS3::PILS3-GFP in the apical hook region (in *pils3-1* background) signal in 3-days-old dark-grown seedlings. Seedlings were grown on solid ½ MS and treated with or without 75 mM NaCl (**A**) or with solvent control DMSO or 5 μg/ml TM (**B**) in liquid ½ MS for 1-4h. Representative images for untreated and 2h time point are shown (see additional images in SFigure 4C, D). Scale bars, 50 μm. n = 10, one-way ANOVA followed by Tukey’s multiple comparison test for each treatment against control or DMSO (* P < 0.05, ** P < 0.01, *** P < 0.001, **** P < 0.0001). **C**, Representative images and relative hypocotyl length of 5-days-old dark-grown seedlings germinated on ½ MS media supplemented with DMSO or 0.15 μg/ml TM. Scale bars, 10 mm, n = 35-50. One-way ANOVA followed by Tukey’s multiple comparison test (b: P < 0.0001). **D**, Relative hypocotyl length of 3-days-old dark-grown seedlings transferred for 2 additional days on ½ MS media supplemented with or without 100 mM NaCl. n = 60, one-way ANOVA followed by Tukey’s multiple comparison test (b: PILS3 OX vs. Col.-0 P < 0.001, PILS3 OX vs. PILS5 OX P < 0.05, PILS3 OX vs *pils2 pils5* P < 0.0001; c: PILS5 OX vs. Col-0 and vs. *pils2 pils5* P < 0.0001; d: *pils2 pils5* vs. Col-0 P < 0.05). **E**, Relative hypocotyl length of 3-days-old dark-grown seedlings transferred for 2 additional days on ½ MS media supplemented with DMSO or 0.5 μg/ml TM. n = 90-150, pooled data of three biological replicates are shown. One-way ANOVA followed by Tukey’s multiple comparison test (b: PILS3^OX^ vs. Col.-0 P < 0.001, PILS5^OX^ vs. Col-0 P < 0.0001; c: *pils2 pils5* vs. Col-0 P < 0.05, *pils2 pils5* vs PILS3 and 5^OX^ P < 0.0001). In all panels boxplots: Box limits represent 25th percentile and 75th percentile; horizontal line represents median. Whiskers display min. to max. values. Representative experiments are shown and all experiments were repeated at least three times.

In agreement with the stabilization of PILS proteins, we observed that chronic ER stress, such as germinating seedlings on TM-containing plates, also strongly enhanced the PILS5-induced growth repression in dark-grown hypocotyls (Figure 4C). To provoke milder ER stresses, we transferred 3 days old dark-grown seedlings for another 2 days to TM containing medium. During these 2 days, the growth of wild-type seedlings was only slightly affected, but PILS5 overexpressing seedlings still showed quantitatively enhanced growth repression (Figure 4D), suggesting that the stabilization of PILS proteins contributes to salt-induced repression of growth rates. Similarly, we also observed hypersensitivity of PILS5 overexpressors when transferred to high salt-containing plates (Figure 4E). In agreement with its effect on protein abundance, also the constitutive expression lines of *PILS3* showed hypersensitivity to ER-stress-inducing conditions (Figure 4D, E), again pointing that the response is not specific to PILS5. This finding is also in agreement with a highly redundant function of *PILS* genes and at least *PILS2* and *PILS5* redundantly control hypocotyl growth in the dark (3). Conversely to the overexpression phenotypes, *pils2 pils5* mutants were less sensitive when transferred to salt or TM when compared to the wild type (Figure 4D, E). This finding illustrates that ER stress signals repress growth in dark-grown hypocotyls at least partially in a PILS-dependent manner.

We noted that the *imp2;* PILS5-GFP^OX^ mutant enhanced PILS5-induced growth repression in dark grown hypocotyls, but not in roots of light grown seedlings (SFigure 1D). This points at a tissue-dependent effect and we hence tested if also the impact of ER-stress on PILS5 proteins is tissue specific. Notably, the induction of ER stress did not increase but lowered the PILS5 abundance in roots (SFigure 5A, B), correlating with PILS5-dependent root growth control (SFigure 5C). Auxin defines plant growth in a concentration and tissue dependent manner, leading to a preferential stimulation and repression of growth in aerial and root tissues, respectively. We accordingly conclude that ER stress differentially affects PILS proteins in shoot and roots and thereby to an overall retardation of growth.

In conclusion, we illustrate that ER-stress perception defines the protein abundance of PILS proteins, which has consequences for auxin signaling rates. We accordingly conclude that the ER stress response machinery utilizes PILS proteins to provoke growth retardation.

## Concluding Remarks

The ER stress response machinery provides a fundamental mechanism to sense and react to environmental stresses. A variety of environmental conditions lead to the accumulation of misfolded proteins or altered composition of membrane lipids in the ER. The imbalance in these biochemical processes is sensed and activates the UPR (28, 29). Defects in the UPR sensor *IRE1* affect auxin signaling output, which may relate to the transcriptional regulation of auxin receptors as well as auxin transport components (30). Here we show that ER stress-inducing conditions define the turnover of PILS proteins and thereby link the fundamental ER stress machinery to auxin-dependent growth control. The here uncovered posttranslational effect on PILS proteins could therefore in part mechanistically explain the interrelation of UPR and auxin signaling.

We uncover that ER stress specifically stabilizes PILS proteins in dark-grown hypocotyl, which consequently represses the nuclear auxin signaling output, leading to growth retardation. The ER stress-dependent control of PILS turnover is tissue-specific, showing reduced and increased PILS turnover in shoot and root tissues. The underlying tissue-specific cues remain to be investigated, but they seem to guide the biphasic auxin responses in shoots and roots, where auxin acts as a promoter and repressor of growth, respectively. It remains however until now completely unknown how PILS turnover is molecularly defined and hence it is difficult to anticipate its tissue specific regulation.

Increasing evidence already suggested that PILS proteins are important players to incorporate environmental signals into developmental growth programs (3–5, 9, 10). Here we show the posttranslational control of PILS protein levels also allows to integrate ER stress-inducing conditions, including imbalanced lipid homeostasis, salt stress, as well as unfolded proteins, with auxin signaling output. We accordingly propose that PILS proteins provide flexibility to adaptive plant development.

In conclusion, our work mechanistically links ER-stress responses to PILS-dependent control of auxin-reliant growth. Accordingly, plant growth retardation under stressful environments is at least in part independent of biochemical limitations and depends on alterations in PILS-dependent auxin signaling output.

## Figure legends

**SFigure 1:** Imbalance in membrane lipids affects PILS-dependent growth

**A**, Sketch of *imp2* mutation in the *CHER1* locus. The change of G to A in *imp2* results in the conversion of glycine (Gly) to arginine (Arg) at the amino acid residue 643. The green boxes represent exons, the blue boxes 5’UTR and 3’UTR. **B**, Representative images and quantification of dark-grown hypocotyls, comparing Col-0 (WT), *cher1-4*, and *imp2*. **C**, Representative images and root length quantification of 5-days-old Col-0 (WT), PILS5-GFP^OX^, *cher1-4, imp2*, and *imp2*; PILS5-GFP^OX^ mutants. n = 20, one-way ANOVA followed by Tukey’s multiple comparison test (distinct letter: P < 0.0001). Scale bars, 100 μm. **D**, Phospholipid analysis of roots from 7-days-old Col-0 wild-type, PILS5-GFP^OX^, *cher1-4* and *imp2* mutants. Lipids are grouped into phosphatidylserine (PS), phosphatidylcholine (PC), phosphatidic acid (PA), phosphatidylethanolamine (PE), phosphatidylinositol (PI) and phosphatidylglycerol (PG). One-way ANOVA followed by Tukey’s multiple comparison test for each phospholipid group (b: P < 0.0001). In all panels with boxplots: Box limits represent 25th percentile and 75th percentile; horizontal line represents median. Whiskers display min. to max. values. Representative experiments are shown and all experiments were repeated at least three times.

**SFigure 2:** *imp2* mutants enhance PILS5

**A, B**, Kinetics of apical hook development in *Col-0* wild-type, PILS5-GFP^OX^ and *imp2;* PILS5-GFP^OX^ germinated on ½ MS media (A) or PILS5^OX^ germinated on ½ MS media supplemented with solvent control DMSO or 0.5 μM FB1 (B). n ≥ 12, statistical significance was evaluated by non-linear regression and a subsequent extra sum of squares F test. End of maintenance phase (*X*_*0*_) and speed of opening (*K*) were compared to *Col-0* wild-type (D) or DMSO control (E) (b, c: P < 0.0001). **C**, Representative images of presumably perinuclear localization of PILS5-GFP in FB1-treated PILS5-GFP^OX^ and *imp2;* PILS5-GFP^OX^ mutants. Scale bars, 50 μm. **D**, qPCR analysis detecting transcript levels of BIP1, BIP2 and PDI6 normalized against UBQ5 and ElF4. Bars represent means ± SD, n = 3. In all panels with boxplots: Box limits represent 25th percentile and 75th percentile; horizontal line represents median. Whiskers display min. to max. values. Representative experiments are shown and all experiments were repeated at least three times.

**SFigure 3:** ER-stress increases PILS protein abundance

**A**, Immunoblot of 4-days-old dark-grown PILS5-GFP^OX^, PILS6-GFP^OX^ and p35S::GFP-HDEL seedlings treated with or without 75 mM NaCl in liquid ½ MS for 1h. α-Actin antibody was used for normalization. **B**, Immunoblot of 4-days-old dark-grown PILS5-GFP^OX^, PILS6-GFP^OX^ and p35S::GFP-HDEL seedlings germinated on ½ MS media with DMSO or 0.15 μg/ml TM. α-Actin antibody was used for normalization. Representative experiments are shown and all experiments were repeated at least three times.

**SFigure 4:** ER-stress exerts a fast effect on PILS proteins

**A-D**, Representative images of p35S::GFP-PILS3 hypocotyls (**A, B**) or pPILS3::PILS3-GFP apical hooks in the *pils3-1* background (**C, D**) in 3-days-old dark-grown seedlings. Seedlings were grown on solid ½ MS and treated with or without 75 mM NaCl (**A, C**) or with solvent control DMSO or 5 μg/ml TM (**B, D**) in liquid ½ MS for 1-4h. Scale bars, 50 μm. Signal intensity was measured (A, B) and statistics is based on a one-way ANOVA followed by Tukey’s multiple comparison test for each treatment against control or DMSO (* P < 0.05, ** P < 0.01, *** P < 0.001, **** P < 0.0001). Representative experiments are shown and all experiments were repeated at least three times.

**SFigure 5:** ER-stress induces PILS5 turnover in roots

**A-B**, Representative confocal microscope images and quantification (A) as well as anti-GFP immunoblot (B) of 4-days-old light grown p35S::PILS5-GFP (PILS5-GFP^OX^) seedlings. Plants were treated with DMSO or 5 μg/ml TM for 6h (A) or 3 hours (B). **C**, 3-days-old light-grown wild-type (WT), p35S::PILS5-GFP (PILS5-GFP^OX^), and *pils2 pils3 pils5* triple mutants (*pils235*) were transferred to TM containing plates (ranging from 25-100 ng/ml) for another 4 days. Relative (to untreated control) root length measurement is shown. Scale bars, 100 μm (A); 0.5cm (C). boxplots: Box limits represent 25th percentile and 75th percentile; horizontal line represents median. Whiskers display min. to max. values. n= 9-11 roots (A). Statistics is based on a t-test (A) or a one-way ANOVA followed by Tukey’s multiple comparison test (C) for each treatment against control (* P < 0.05, ** P < 0.01, *** P < 0.001, **** P < 0.0001). Error bars ± SD, n = 29-32 seedlings (C). Representative experiments are shown and all experiments were repeated at least three times.

## Material and methods

### Plant material and growth conditions

Arabidopsis thaliana Col-0 (wild type), *p35S::PILS5-GFP* (3), *p35S::GFP-PILS3* (4), *p35S::PILS6-GFP* (5), pPILS3::PILS3-GFP (4), pDR5::GFP (32), *pils2 pils5* (3), *pils2 pils3 pils5* (9), *cher1-4* (33), pCHER1::CHER1-YFP (11), p35S::GFP-HDEL (34), *p35S::DER1-mScarlet* (10). Seeds were stratified at 4°C for 2 days in the dark. Seedlings were grown vertically on half Murashige and Skoog medium (1/2 MS salts (Duchefa), pH 5.9, 1% sucrose, and 0.8% agar). Plants were grown under long-day (16 h light/8 h dark) or under dark conditions at 20–22°C.

### EMS mutagenesis, forward genetic screen, and sequencing

The EMS screen for *imperial PILS* (*imp*) mutants has been described previously (9). Firstly, *imp2* was mapped on the chromosome 3 between T21E2MspI (4.981 Mb) and MSJ11 (5.315 Mb). Then, 170 individuals of F2 progeny derived from cross of *imp2* with *Col-0* were selected based on the dark-grown hypocotyl phenotype. The selected seedlings were transferred to soil. For next generation sequencing the genomic DNA of *imp2* was isolated using the DNeasy Plant Mini Kit (Qiagen), according to the manufacturer’s instructions. The DNA samples were sent to BGI Tech (https://www.bgi.com) for whole genome re-sequencing using Illumina’s HiSeq 2000.

### Kinetics of apical hook development

Seedlings were grown in a light protected box equipped with an infrared light source (880 nm LED) and a spectrum-enhanced camera (EOS035 Canon Rebel T3i) modified by Hutech technologies with a built-in, clear, wideband-multicoated filter. The camera was operated by EOS utility software. Angles between the cotyledons and the hypocotyl axis were measured every 3 h in the dark until opening using ImageJ (http://rsb.info.nih.gov/ij/) software. The complementary angle of the measured angle is reported in the graphs (180° represents full closure and 0° full opening). More information can be found in (4).

### Chemicals and Treatments

Tunicamycin (TM) (Santa Cruz) and fumonisin B1 (FB1) (Santa Cruz) were all dissolved in DMSO (Duchefa). NaCl was added directly to the medium. Treatments with TM and FB1 were performed on 3-4-day old dark grown seedlings (transferred to supplemented media) or germinated directly on the respective compound.

### Phospholipid Analysis

Arabidopsis roots (around 200-300 mg fresh weight) were collected from vertical agar plates, weighted and immediately transferred into glass tubes containing 1 ml of isopropanol, the samples were treated at 80°C for 5 min to inactivate phospholipase activities. Lipids were extracted with methyl-*tert*-butyl ether (MTBE) Methanol/H_2_O (100:30:4, v/v/v) solvent mix (38). Phospholipids separation was performed on Merck HPTLC silica gel 60 (20 x 10 cm) with the following migration solvent: CHCl_3_/Methanol/2-propanol/KCl (0.25% w/v in water)/methylacetate/trimethylamine 15/5/12.5/4/12/1.5, v/v/v/v/v/v). Lipids were visualized by spraying 2 mg/ml (in acetone/water 8/2, v/v) on plates. After drying, HPTLC plates were imaged with a ChemiDoc (BioRad). Lipid bands were scratched from the plates and their fatty acids extracted (fatty acid methyl esters FAMEs) and quantified by GC-MS (Agilent 7890 A and MSD 5975 Agilent EI) as in (39). After normalization to the lipid standard C17:0 and to the fresh weight, the values obtained were expressed in nmol of fatty acids mg^-1^ FW. The value for each lipid class is the sum of all fatty acids found in this class and is an average of 3 biological replicates.

### RNA Isolation and qPCR

RNA was isolated using inuPREP Plant RNA Kit (Analytic Jena) following manufactures instructions. qPCR has been performed as described in Feraru et al., 2019. The primers are listed in a STable 1.

### Microscopy

Confocal microscopy was done with a Leica SP8 (Leica). Fluorescence signals for GFP (excitation 488 nm, emission peak 509 nm), mScarlet-i (excitation 561 nm, emission peak 607 nm) and YFP (excitation 513 nm, emission peak 527 nm) were detected with a 10x or 20x (dry and water immersion, respectively) objective. Z-stacks were recorded with a step size of 840 nm. On average, 24 slices were captured, resulting in an average thickness of approximately 20 μm. Image processing was performed using LAS AF lite software (Leica).

### Protein Extraction and Immunoblot (IB) Analysis

Seedlings were ground to fine powder in liquid nitrogen and solubilized with extraction buffer (25 mM TRIS, pH 7.5, 10 mM MgCl_2_, 15 mM EGTA, 75 mM NaCl, 1 mM DTT, 0.1% Tween20, with freshly added proteinase inhibitor cocktail (Roche). After spinning down for 60 min at 4°C with 20.000 rpm supernatant was transferred to a new tube and the protein concentration was assessed using the Bradford method. Protein extracts were used for immunoblot with anti-GFP (Roche #11814460001, 1:1,000), anti-RFP (Chromotek #6g6, 1:1,000) or anti-Actin (Sigma #A0480, 1:10,000) and goat anti-mouse IgG (Jackson ImmunoResearch # 115-036-003, 1:10,000) for detection.

### Statistical analysis and reproducibility

GraphPad Prism software 9 was used to evaluate the statistical significance of the differences observed between control and treated groups and to generate the graphs. All experiments were, if not stated different, always repeated at least three times and the depicted data show the results from one representative experiment.

## Acknowledgments

We are grateful to Ykä Helariutta for sharing published material; our team members for helpful discussions; to the Bordeaux Metabolome Facility MetaboHUB (ANR-11-INBS-0010) and the LIC Imaging Center Freiburg for expertise and support. This work was supported by Vienna Research Group (VRG) program of the Vienna Science and Technology Fund (WWTF to J.K-V.), the Austrian Science Fund (FWF) (P29754 to J.K-V., P33497 to S.W. and Hertha Firnberg T728-B16 and Elise Richter V690-B25 to E.F.), the European Research Council (ERC) (639478-AuxinER to J.K-V.), German Science fund (DFG; 470007283 and CIBSS – EXC-2189 to J.K.-V.), Austrian Academy of Sciences (25479 to J.F.) and the French National Research Agency (ANR-18-CE13-0025 to Y.B.).

## Author contributions

S.W., J.F., E.F., C.B., M.I.F, L.M.-M. and L.S. performed most of the experiments. Y.B. conducted the phospholipid analysis. S.W. and J.K.-V. devised and coordinated the project and wrote the manuscript. All authors saw and commented on the manuscript.

