## Supplemental Table 1 for "Endoplasmic-Reticulum stress controls PIN-LIKES abundance and thereby growth adaptation"

| Gene | Primer name | Primer sequence | purpose |
|---|---|---|---|
| BIP1/2 | BIP1/2_qPCR_FW | CCACCGGCCCCAAGAG | qPCR |
| BIP1/2 | BIP1/2_qPCR_REV | GGCGTCCACTTCGAATGTG | qPCR |
| PDI6 | PDI6_qPCR_FW | CGAAGTGGCTTTGTCATTCCA | qPCR |
| PDI6 | PDI6_qPCR_REV | GCGGTTGCGTCCAATTTT | qPCR |
| CNX1 | CNX1_qPCR_FW | GTGTCCTCGTCGCCATTGT | qPCR |
| CNX1 | CNX1_qPCR_REV | TTGCCACCAAAGATAAGCTTGA | qPCR |
